# Cytoplasmic pressure maintains epithelial integrity and inhibits cell motility

**DOI:** 10.1101/2021.04.05.438452

**Authors:** Pragati Chengappa, Tia M. Jones, James M. Cowan, Devneet Kainth, Ryan J. Petrie

## Abstract

Cytoplasmic pressure, a function of actomyosin contractility and water flow, can regulate cellular morphology and dynamics. In mesenchymal cells, cytoplasmic pressure powers cell protrusion through physiological three-dimensional extracellular matrices. However, the role of intracellular pressure in epithelial cells is relatively unclear. Here we find that high cytoplasmic pressure is necessary to maintain barrier function, one of the hallmarks of epithelial homeostasis. Further, our data show that decreased cytoplasmic pressure facilitates lamellipodia formation during the epithelial to mesenchymal transition (EMT). Critically, activation of the actin nucleating protein Arp2/3 is required for the reduction in cytoplasmic pressure and lamellipodia formation in response to treatment with hepatocyte growth factor (HGF) to induce EMT. Thus, elevated cytoplasmic pressure functions to maintain epithelial tissue integrity, while reduced cytoplasmic pressure triggers lamellipodia formation and motility during HGF-dependent EMT.

## Introduction

Epithelial cells can become highly motile through the epithelial to mesenchymal transition (EMT). During EMT, cell-cell adhesion can be reduced within an epithelial monolayer to increase the motility of cell clusters and single cells. The small GTPase RhoA and non-muscle myosin II (NMII) are critical regulators of cell-cell adhesion and EMT. Specifically, the formation and maintenance of E-cadherin containing cell-cell junctions and ZO-1-based tight junctions are governed by RhoA signaling and NMII activity through changes in actomyosin contractility (1–4). During EMT, epithelial cells become more mesenchymal in character by changing gene expression, disassembling cell-cell junctions, developing front-back polarity, and forming lamellipodia to increase motility (5–9). Interestingly, inhibiting RhoA signaling or actomyosin contractility can lead to increased cell motility and invasion in multiple cell types, including Madin-Darby canine kidney (MDCK) cells (10–14).

NMII-driven actomyosin contractility also governs cytoplasmic hydraulic pressure in epithelial and mesenchymal cells (15–20). In migrating mesenchymal cells, the magnitude of intracellular pressure can help dictate the types of protrusions that are formed at the leading edge. For example, in fibroblasts migrating through highly cross-linked three-dimensional (3D) matrices, the small GTPase RhoA activates actomyosin contractility to increase pressure and switch cells from actin polymerization-driven lamellipodial protrusions to pressure-based lobopodial protrusions (16,18,21). In proliferating epithelial cells, cell rounding is achieved by the simultaneous increase in cortical actomyosin contractility and water influx to increase intracellular pressure and maintain mitotic cell shape (19,20). Other studies have shown that the epithelial barrier and transport functions can change in response to extracellular osmotic and hydrostatic pressures; reviewed in (22). Critically, the function of intracellular pressure within epithelial cells during interphase remains unclear.

In this study, we tested the hypothesis that actomyosin contractility increases cytoplasmic hydraulic pressure in epithelial cells to prevent lamellipodia formation and cell motility. We find that epithelial cells maintain significantly higher intracellular pressures than fibroblasts and this pressure helps to maintain the integrity of the epithelial monolayer. Further, inducing EMT by treating cells with hepatocyte growth factor (HGF) causes a decrease in cytoplasmic pressure that is required for the formation of lamellipodial protrusions and increased cell migration.

## Results

### High cytoplasmic pressure corresponds with high electrical resistance across epithelial monolayers

To understand whether intracellular pressure could play a role in epithelial cell function, we first compared the intracellular cytoplasmic pressures of primary human dermal fibroblasts and MDCK I cells. Epithelial MDCK I cells had a significantly higher cytoplasmic pressure than the mesenchymal dermal fibroblasts (approximately 3-fold) (Fig. 1 A). Interestingly, MDCK I cell monolayers are reported to have a high electrical resistance compared to MDCK II cells (see (23) for a detailed explanation of the MDCK cell lines used in this report) despite a lack of measurable differences in junction morphology or ZO-1 and E-cadherin protein expression (24). We then hypothesized that the high cytoplasmic pressure in MDCK I cells could help to increase the electrical resistance of epithelial cell monolayers. To test this hypothesis, we first confirmed the published differences in transepithelial electrical resistance (TEER) of MDCK I and II monolayers (Fig. 1 B). Additionally, we determined there were no significant differences in ZO-1 localization to tight junctions between the two cell lines (Fig. 1 C and D) as reported previously (24). Despite the similarities in morphology and ZO-1 protein expression, we found that cytoplasmic pressure was approximately 3-fold lower in MDCK II cells compared to the MDCK I cell line (Fig. 1 E), suggesting there could be a role for cytoplasmic pressure in maintaining the integrity of epithelial monolayers. We tested this hypothesis using the parental MDCK epithelial cell line (23) by experimentally reducing intracellular pressure and measuring the change in TEER. Specifically, MDCK cells were either untreated (control) or treated with hyper-osmotic media for 1 hour before independently measuring the intracellular pressure and the electrical resistance of the monolayer. We found that treating MDCK cells with hyper-osmotic media significantly reduced intracellular pressure (Fig. 1 F), and this change in pressure corresponded with a significant drop in the resistance of the monolayer (Fig. 1 G). Together, these findings show that high cytoplasmic pressure can promote epithelial integrity by preventing the flow of small molecules and ions past tight junctions to maintain the electrical resistance of the monolayer.

**Figure 1.**
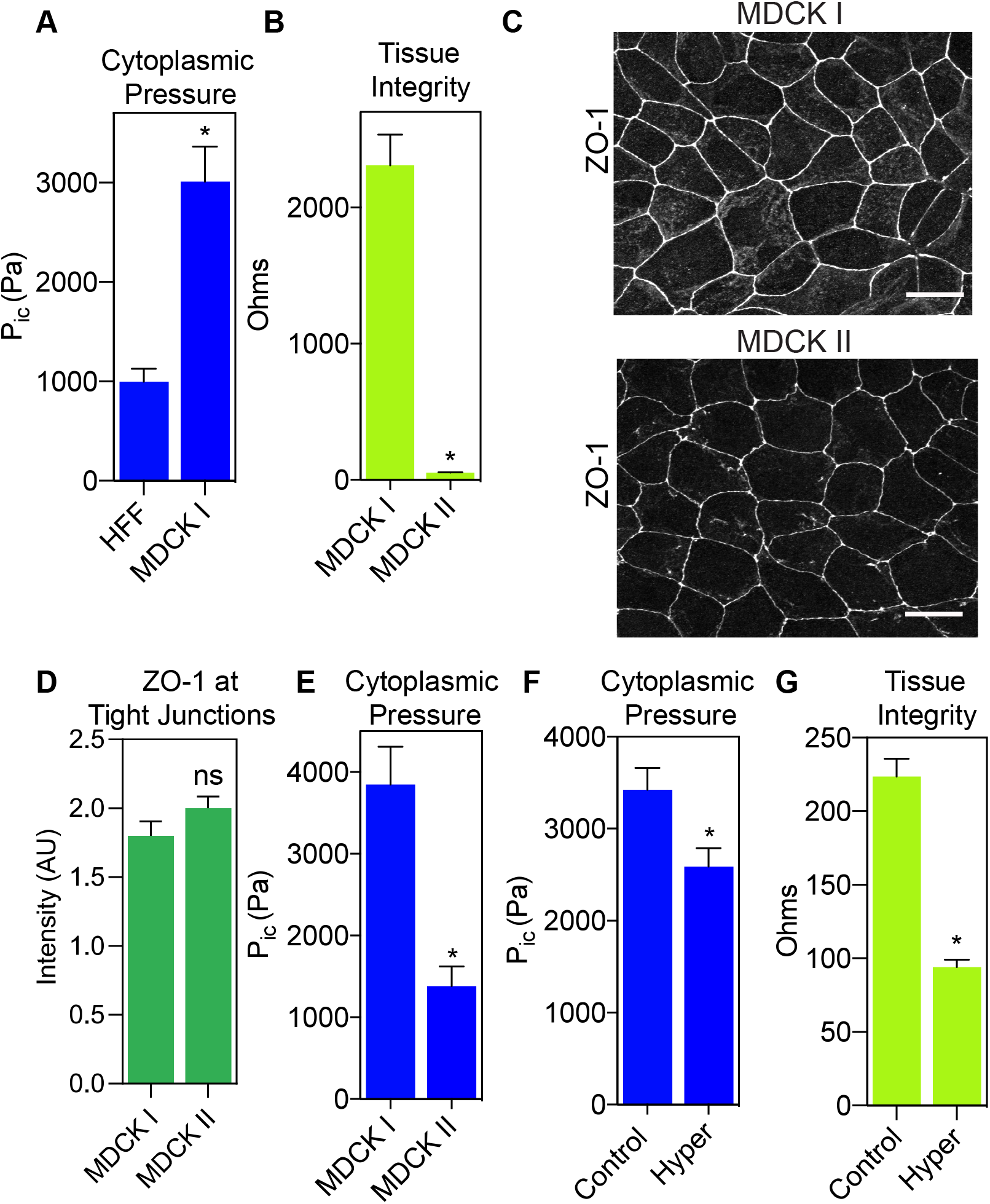
Cytoplasmic pressure helps to maintain the barrier function of epithelial cell monolayers. (A) Epithelial MDCK cells have significantly higher cytoplasmic pressure compared to mesenchymal HFFs (*n* ≥ 29, N = 3). *, P < 0.0001. (B) The MDCK II cell line has significantly lower TEER compared to the MDCK I cell line (*n* ≥ 12, N = 3). *, P < 0.0001. (C) Differences in TEER between MDCK I and MDCK II cell lines do not appear to be due to differences in ZO-1 (greyscale) localization at cell-cell junctions. Bars = 10μm. ZO-1 intensity quantified in (D) (*n* ≥ 30, N = 3). Not significant (ns), P = 0.1412. (E) MDCK I cells have significantly higher intracellular compared to MDCK II cells (*n* ≥ 30, N = 3). *, P < 0.0001. (F) Treatment of parental MDCK cells with hyper-osmotic media significantly lowers intracellular pressure due to flow of water out of the cell (*n* ≥ 30, N = 3). *, P = 0.008. (G) Reducing cytoplasmic pressure in parental MDCK cells increases the permeability of the epithelial monolayer (*n* = 9, N = 3). *, P < 0.0001.

### Decreasing epithelial cytoplasmic pressure increases lamellipodia formation and cell motility

Since low intracellular pressure is associated with lamellipodia formation in migrating fibroblasts (17), we next investigated if triggering epithelial cell motility similarly reduced intracellular pressure. We treated MDCK cells (note that parental MDCK cells are used in the remainder of this report) with HGF to increase epithelial cell motility (9,25). As expected, treatment with HGF increased cell movement and scattering compared to control cells (Fig. 2 A and supplemental movie 1). Further, the HGF-induced motility was associated with significant cortactin-positive lamellipodia formation after only 60 minutes of HGF treatment (Fig. 2 B and C), consistent with a previous report (8). Strikingly, this increase in MDCK cell motility and lamellipodia formation was accompanied by a significant decrease in cytoplasmic pressure compared to control cells (Fig. 2 D). Since HGF treatment is known to induce EMT in epithelial cell monolayers (26,27), we determined whether these changes to protrusion formation, cytoplasmic pressure, and cell movement were accompanied by changes in E-cadherin and vimentin expression, two hallmarks of EMT. We measured no significant changes in E-cadherin or vimentin expression following treatment with HGF for 60 min. compared to control cells (supplemental figure 1 A and B). These results suggest that the drop in cytoplasmic pressure and increase in lamellipodia formation and cell migration preceded EMT-associated changes in protein expression. Together, these data indicate that HGF-induced changes in cytoplasmic pressure and cell dynamics occurs before the induction of a complete EMT in MDCK cells.

**Figure 2.**
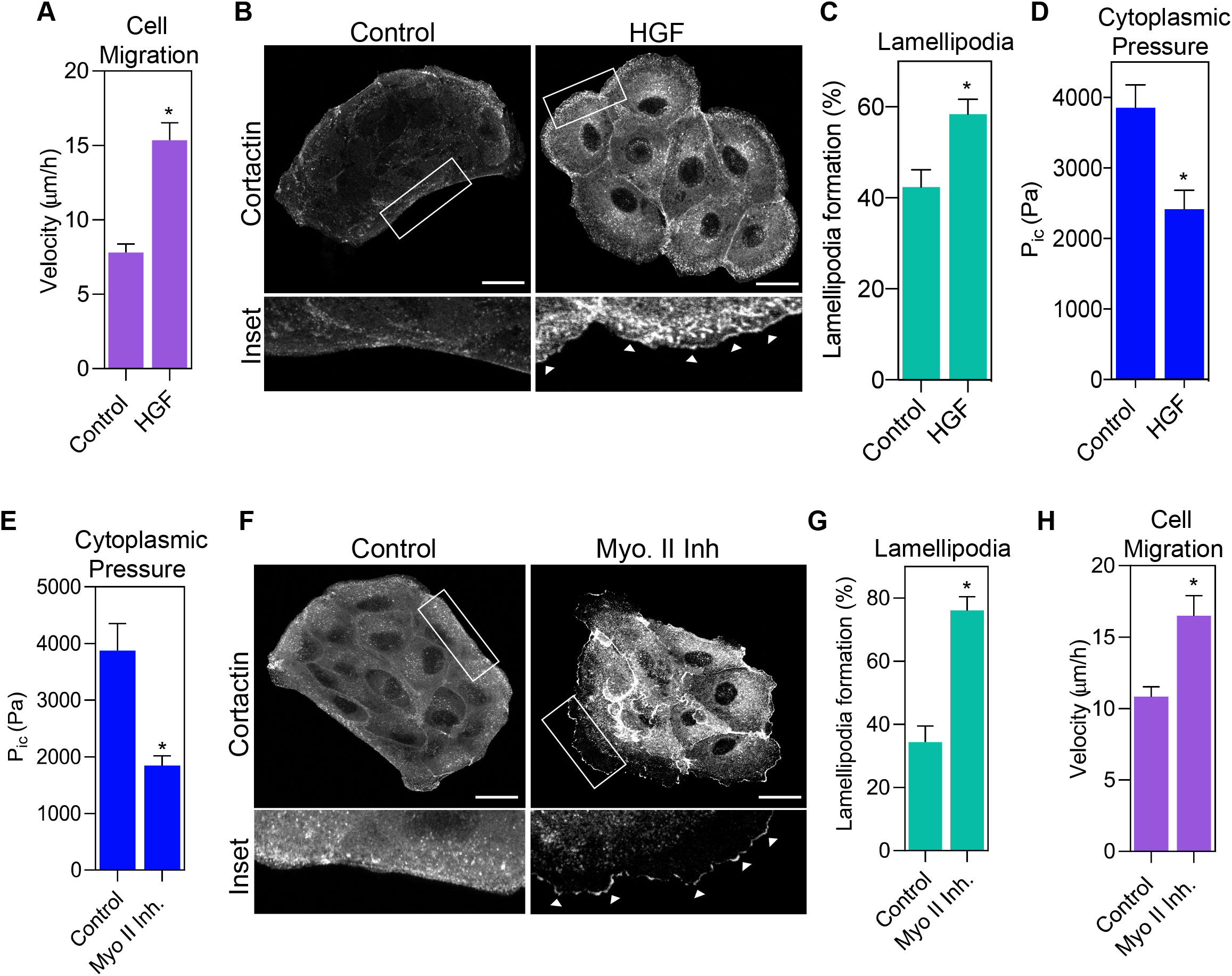
Decreasing pressure through HGF treatment or NMII inhibition increases lamellipodia formation and cell migration. (A) Treatment of parental MDCK cells with HGF caused significantly increased velocity (*n* ≥ 30, N = 3). *, P < 0.0001. (B) Treatment of MDCK cells with HGF significantly increased lamellipodia formation as shown by increased localization of cortactin (greyscale) to the leading edge (white arrowheads). Bars = 10μm. Quantified in (C) (*n* ≥ 30, N = 3). *, P = 0.002. (D) Cells treated with HGF for 1 hour have significantly lower cytoplasmic pressure compared to control cells (*n* ≥ 60, N = 3). *, P = 0.0009. (E) Treatment with the myosin II inhibitor, blebbistatin, significantly decreases intracellular pressure following a 1 hour treatment (*n* ≥ 30, N = 3). *, P= 0.016. (F) Treatment of parental MDCK cells with blebbistatin increased lamellipodia formation indicated by the enrichment of cortactin (greyscale) at the leading edge (white arrowheads) compared to control cells. Bars = 10μm. Quantified in (G) (*n* ≥ 30, N = 3). *, P<0.0001. (H) Myosin II inhibition results in increased cell velocity relative to control cells (*n* ≥ 30, N=3). *, P= 0.023.

Actomyosin contractility generated by non-muscle myosin II (NMII) is essential for controlling MDCK cell scattering (10) and cytoplasmic pressure in fibroblasts (18). To further test the hypothesis that cytoplasmic pressure controls epithelial cell dynamics, we treated cells with the NMII inhibitor blebbistatin and measured changes to cytoplasmic pressure, lamellipodia formation, and cell migration. Treatment of MDCK cells with 25 μM blebbistatin for 60 min significantly decreased intracellular pressure (Fig. 2 E), which is similar to its effect on pressure in high-pressure fibroblasts migrating through 3D matrices. Critically, this decrease in epithelial cytoplasmic pressure corresponded with significant increases in lamellipodia formation and cell velocity (Fig. 2 F-H, and supplemental movie 2). Thus, NMII activity is required to maintain the high cytoplasmic pressure in MDCK cells. Further, reducing pressure by directly inhibiting NMII is sufficient to increase lamellipodia formation and cell motility.

### Cell-cell adhesion is not required for generation of cytoplasmic pressure in epithelial cells

Since NMII activity generates cytoplasmic pressure in epithelial cells and cell-cell junctions are important regulators of NMII activity (2,4), we tested the hypothesis that Ca^2+^-dependent cell-cell junctions are required to increase NMII activity to enhance contractility and prevent cell movement. Cells were treated with 3mM EGTA for 60 minutes at 37 °C to chelate Ca^2+^ ions and disrupt E-cadherin interactions between adjacent cells, as previously reported (28). The loss of cell-cell junctions was confirmed by measuring the extent of colocalization between F-actin and E-cadherin, as well as qualitatively by the formation of visible gaps between adjacent cells (Fig. 3 A and B). HGF treatment decreased cytoplasmic pressure before any pronounced cell scattering was detected (Fig. 3 B and C; pressures are normalized to control values due to variability in baseline pressures). Strikingly, disruption of cell-cell adhesions following EGTA treatment did not significantly affect intracellular pressure (Fig. 3 C) or cell velocity (Fig. 3 D). Taken together, these findings show that Ca^2+^-dependent cell-cell junctions are not required to generate cytoplasmic pressure in epithelial cells. Further, changes in intracellular pressure as a result of HGF treatment occur before large-scale remodeling of E-cadherin containing junctions.

**Figure 3.**
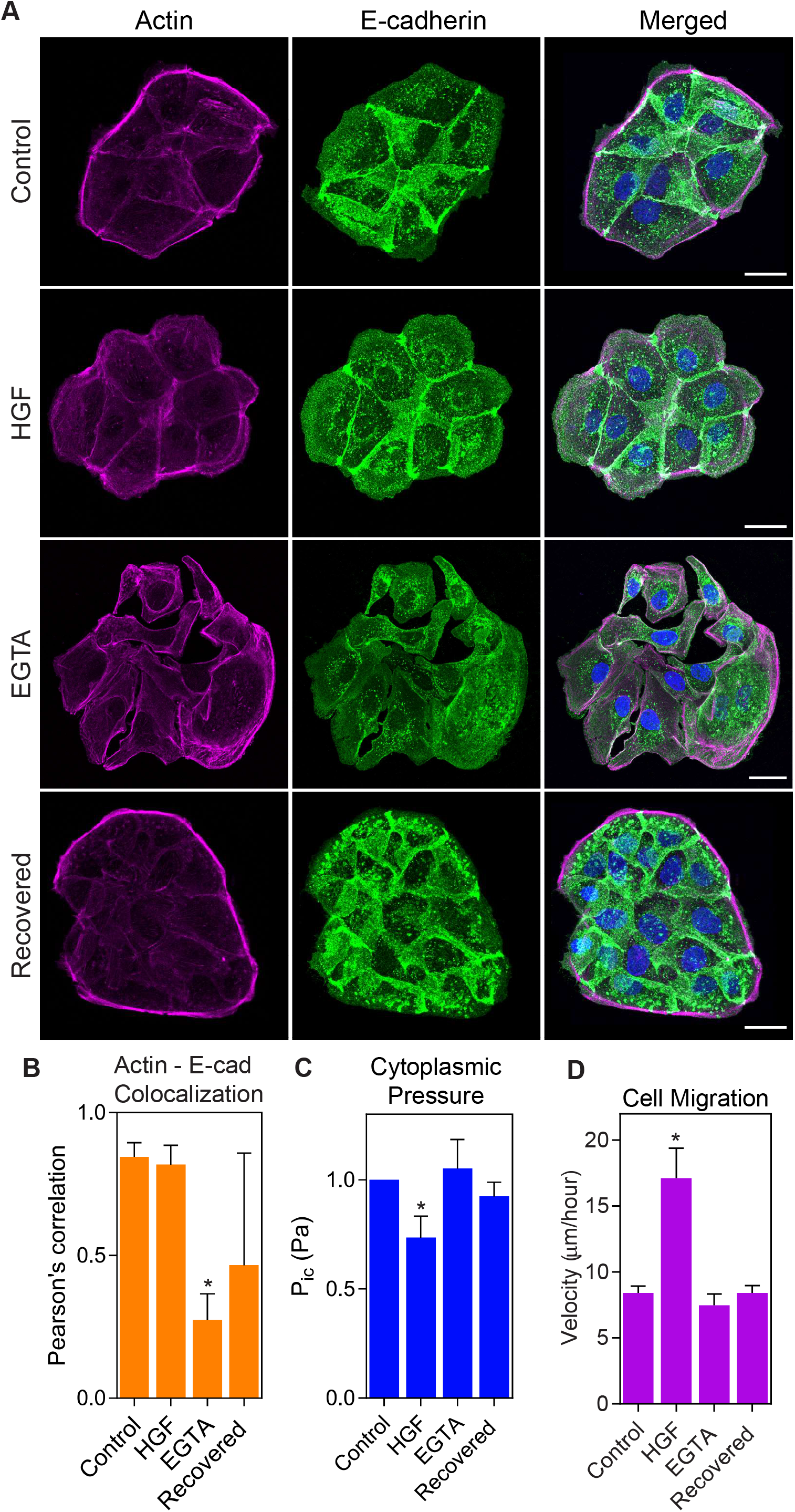
Cytoplasmic pressure is not regulated by calcium-dependent cell-cell adhesion. (A) Control, HGF-treated, EGTA-treated, and recovered cells were stained for E-cadherin (green) and F-actin (magenta) to assess cell-cell adhesion. Bars = 10μm. (B) E-cadherin-actin colocalization was significantly lower in EGTA treated cells compared to control cells, indicating reduced cell-cell adhesion (*n* ≥ 30, N = 3). *, P = 0.034 versus control. (C) HGF treatment reduced intracellular compared to control cells, while there was no change in pressure following EGTA treatment or recovery (*n* ≥ 30, N = 3). *, P = 0.013 versus control. (D) HGF treatment significantly increased cell motility compared to control cells, while EGTA treatment and recovery caused no significant change to cell movement (*n* ≥ 30, N = 3). *, P < 0.0001 versus control.

### Increasing osmotic pressure prevents HGF-triggered motility

Since HGF treatment decreased cytoplasmic pressure and increased lamellipodia formation and cell velocity (Fig. 2), we next tested whether maintaining high intracellular pressure would prevent HGF-triggered changes to protrusion formation and cell migration. HGF treatment significantly reduced cytoplasmic pressure (Fig. 4 A), as expected. Critically, treating cells with HGF in hypotonic (hypo) media (complete media diluted 1:5 with H_2_O) prevented HGF treatment from decreasing intracellular pressure. This hypotonic media was selected based on its ability to increase intracellular pressure in primary fibroblasts (17). Further, MDCK cells were unable to increase the formation of cortactin-positive lamellipodial protrusions when treated with HGF in hypotonic media compared to treatment with HGF alone (Fig. 4 B and C). This reduced lamellipodia formation corresponded to a significant decrease in cell velocity (Fig. 4 D). Specifically, time-lapse brightfield imaging of live cells co-treated with hypotonic media and HGF revealed-significantly decreased velocity compared to cells treated with just HGF. These results suggest that the reduction of intracellular pressure is required for the formation of lamellipodia and the increase in cell motility following HGF treatment.

**Figure 4.**
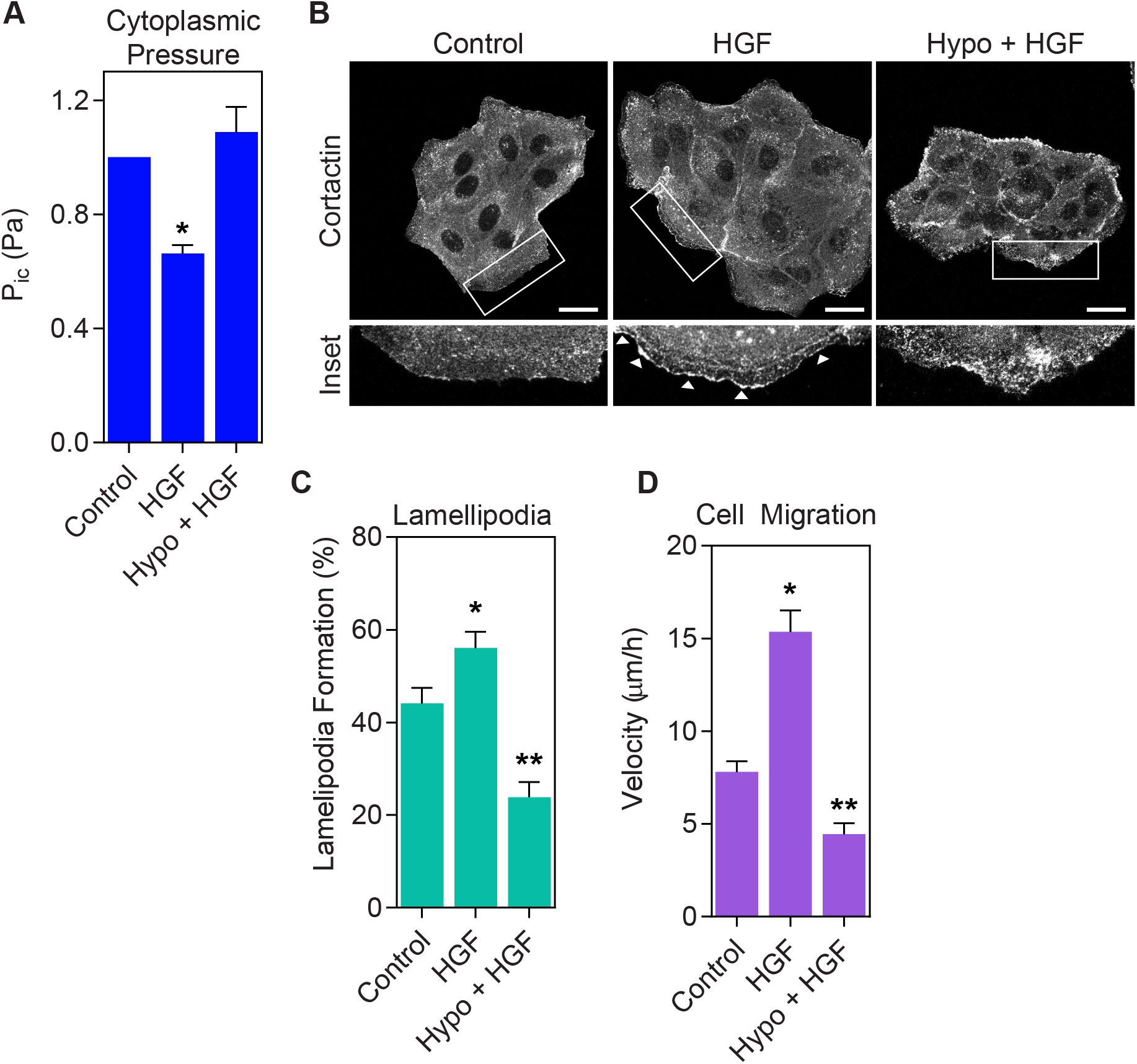
Preventing cytoplasmic pressure from decreasing inhibits lamellipodia formation and cell migration in response to HGF. (A) Parental MDCK cells treated 70% hypo-osmotic media for 1 hour prevents HGF treatment from reducing cytoplasmic pressure (*n* ≥ 45, N = 3). *, P < 0.02 versus control. (B) Representative images of MDCK cells treated as indicated and immunostained for cortactin (greyscale) to assess lamellipodia formation (white arrowheads). Bar = 20μm. Quantified in (C) (*n* ≥ 35, N = 3). *, P < 0.04 versus control. **, P < 0.0001 versus HGF. (D) Treating parental MDCK cells with hypo-osmotic media prevents HGF from increasing cell motility (*n* ≥ 30, N = 3). *, P < 0.0001 versus control. **, P < 0.0001 versus HGF.

### Arp2/3 activity is required to reduce pressure and increase motility in response to HGF

Given the clear link between reduced pressure, lamellipodia formation, and increased cell motility, we tested the hypothesis that low-pressure lamellipodia formation is essential for the movement of epithelial cells following HGF treatment. We predicted that preventing lamellipodia formation by inhibiting the actin nucleating protein Arp2/3 would inhibit the HGF-dependent increase in cell velocity. Inhibition of Arp2/3 with a combination of CK666 and CK869 for 60 min (29) did not affect lamellipodia formation or cytoplasmic pressure compared to control cells (Fig. 5 A-C). However, treating cells with a combination of HGF and the Arp2/3 inhibitors maintained high cytoplasmic pressure (Fig. 5 C), prevented lamellipodia formation (Fig. 5 A and B), and significantly inhibited cell motility (Fig. 5 D). Together, these data suggest that Arp2/3 activity is required for decreasing intracellular pressure and increasing cell motility following HGF treatment.

**Figure 5.**
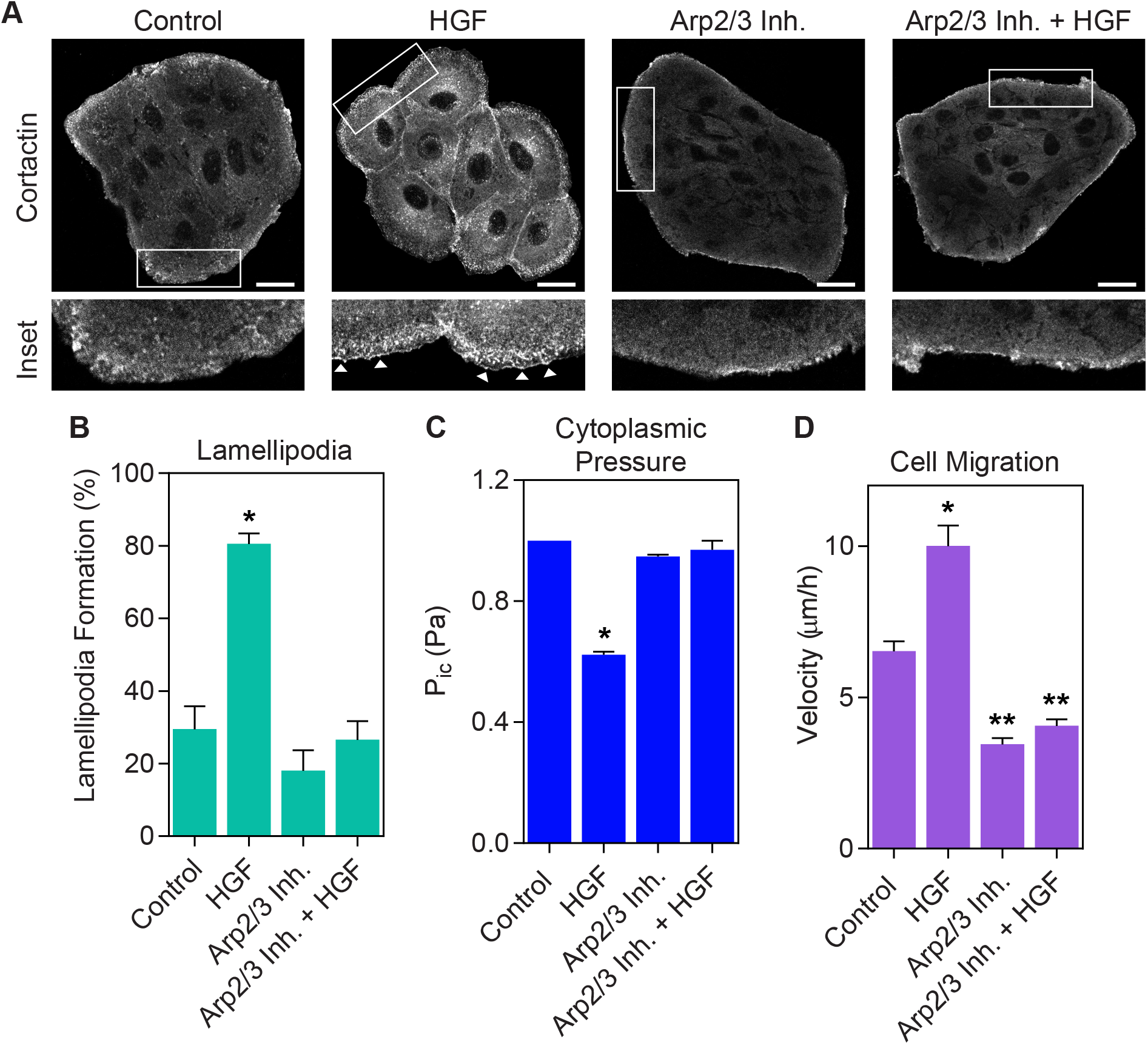
Arp2/3 activity is required to reduce cytoplasmic pressure and increase epithelial cell migration. (A) Representative images of parental MDCK cells co-treated with HGF and the Arp2/3 inhibitors CK666 and CK869, immunostained for cortactin (greyscale). Inhibition of Arp2/3 prevents cortactin-positive lamellipodia from forming (white arrowheads). Bar = 20μm. Quantified in (B) (*n* ≥ 15, N = 3). *, P < 0.0001 versus control. (C) Inhibiting Arp2/3 activity prevents HGF treatment from reducing the cytoplasmic pressure of parental MDCK cells (*n* ≥ 30, N = 3). *, P < 0.0001 versus control. (D) Inhibition of Arp2/3 activity prevents HGF treatment from increasing MDCK cell motility (*n* ≥ 30, N = 3). *, P < 0.0001 versus control. **, P < 0.0001 versus HGF.

## Discussion

Epithelial cells maintain a significantly higher cytoplasmic pressure compared to fibroblasts. This high pressure appears to have at least two distinct functions in epithelial cells. First, cytoplasmic pressure promotes tissue integrity which prevents small molecules from passing through a confluent epithelial monolayer. This barrier function is consistent with the established use of hyper-osmotic fluid to transiently open the blood-brain barrier and facilitate the passage of small molecules (30,31). Second, high intracellular pressure is critical in maintaining epithelial cells in a non-motile state. Initiating cell motility by treating cells with HGF decreases cytoplasmic pressure. This drop in pressure is required for subsequent lamellipodia formation, increased cell migration, and monolayer dissolution. Mechanistically, epithelial cell pressure is governed by NMII activity, similar to mesenchymal cells (32), and this activity is not a result of cell-cell adhesion. Importantly, Arp2/3 activity is required for the decrease in pressure in response to HGF treatment, and this is consistent with its negative regulation of hydraulic pressure in fibroblasts (17,33). Overall, intracellular pressure and its regulation represents a new physical mechanotransduction mechanism that will help us to better understand the established role of actomyosin contractility in regulating EMT and epithelial cell dynamics.

## Materials and Methods

### Reagents and cell culture

The following reagents were used in this study: blebbistatin (Thermo Fisher Scientific) used at 25 μM, HGF (R&D Systems) used at 25 ng/ml, CK666 (Sigma) used at 200 μM, and CK869 (Sigma) used at 30 μM. Cells were treated with the indicated inhibitors for 1-2 h at 37°C. For hypo-osmotic media, regular media was diluted 1:5 with H2O, as previously reported (17). For hyperosmotic solutions, 300 mM sorbitol (MilliporeSigma) was applied.

MDCK I and II cells (23,24) are maintained by the European Collection of Authenticated Cell Cultures (ECACC) and were purchased through MilliporeSigma. Human foreskin fibroblasts (HFFs) and parental MDCK (NBL-2) cells were purchased from the American Tissue Type Collection (ATCC). All cells were used at passages 8-25 and were maintained in phenol red-free DMEM (HyClone) containing 7.5% FBS (MilliporeSigma), 4.5 g/liter glucose, 100 U/ml penicillin, 100 mg/ml streptomycin (Invitrogen), and 2mM L-glutamine (Life Technologies) at 37°C and 10% CO_2_.

### Antibodies and immunofluorescence imaging

Cells were fixed with 4% paraformaldehyde (Electron Microscopy Science), permeabilized with 0.25% Triton X-100, and blocked with 0.2% BSA in PBS. All antibodies were diluted in 0.2% BSA before applying to cells. The following antibodies were used for immunofluorescence: mouse anti-cortactin (EMD Millipore), rabbit ZO-1 (Invitrogen), mouse anti-E-cadherin (BD Biosciences), goat anti-mouse IgG Alexa 488 (Invitrogen), goat anti-rabbit IgG Alexa 568 (Invitrogen), goat anti-rabbit IgG Alexa 647 (Invitrogen), rhodamine phalloidin (Invitrogen), and DAPI (Thermo Fisher Scientific). Cells were imaged using a scanning confocal microscope with multi-line Argon (458 nm, 488 nm, or 515nm) lasers (LSM 700-Karl Zeiss or IX83-Olympus

FluoView) with a 63X, 1.4 NA oil objective or an IncuCyte S3 Live-Cell Analysis System (Sartorius) with a 10X objective. Brightness and contrast were linearly adjusted using ImageJ (National Institutes of Health). The following antibodies were used for western blotting: mouse anti-E-cadherin (BD Biosciences), mouse anti-vimentin (Millipore Sigma), and mouse anti-GAPDH (Fitzgerald Industries International). Colocalization was measured using the Colocalization plugin on Fiji (National Institutes of Health). The actin and E-cadherin channels were separated, and a region of interest was drawn around the cell in Channel 1 or the actin channel. These images were thresholded using the Costes regression threshold command in the Coloc2 plugin which automatically thresholds the images by identifying the maximum intensity threshold of each channel. The number of overlapping pixels between the actin Channel 1 and E-cadherin Channel 2 were measured. This measurement represents the colocalization between actin and E-cadherin in the cells measured. A lower Pearson’s correlation value indicates less correlation between actin and E-cadherin, which correlates with decreased junctional integrity.

### Transepithelial electrical resistance (TEER)

TEER, a measure of monolayer integrity and tight-junction permeability, was measured by seeding MDCKs into a 12 mm trans-well insert with 0.4mm pore size (Corning) and allowing them to form a monolayer for 72 h. After treatment, TEER was measured using an EVOM2 Voltohmmeter (WPI) and EndOhm chamber (WPI). TEER values obtained in the absence of cells (trans-well insert) were considered as background and were subtracted from the total TEER (cell monolayer plus insert). Resistances were calculated in ohms. All TEER measurements were performed in an environmental chamber maintained at 37°C and 10% CO_2_.

### Motility assays

MDCK cells were seeded in 7.5% FBS in DMEM onto 35mm glass bottom dishes (WPI). Three days after seeding, time-lapse movies were captured in chambers set at 10% CO_2_ and 37°C using a 32x, 0.4 NA Ph1 objective on a 510 NLO mETA AxioObserver Z1 (Zeiss). Cells were tracked every 5 or 10 min for 12 h using the Manual Tracking plugin (F. Cordelieres, Institute Curie, Paris, France) with ImageJ. Velocity was calculated from the tracking data using the ImageJ Chemotaxis and Migration Tool plugin (Ibidi). The measured velocities were plotted with Prism 7 (GraphPad).

### Intracellular pressure measurements

Direct measurements of intracellular pressure were performed using the 900A micropressure system (WPI) according to the manufacturer’s instructions and as described (34). Briefly, a 0.5 μm micropipette (WPI) was filled with 1 M KCl solution, placed in a microelectrode holder half-cell (WPI), and connected to a pressure source regulated by the 900A system. A calibration chamber (WPI) was filled with 0.1 M KCl and connected to the 900A system, and the resistance of each microelectrode was set to zero and then secured in an MPC-325 micromanipulator (Sutter Instrument Company) within an environmental chamber maintained at 37°C and 10% CO_2_, on an LSM 700 microscope (Carl Zeiss). To measure intracellular pressure, the microelectrode was driven at a 45° angle into the cytoplasm equidistant between the nucleus and the plasma membrane, maintained in place for ~5 sec, and removed. The pressure measurement was calculated as the average pressure reading during this interval of time.

### Calcium switch assay

Ethylene glycol-bis(2-aminoethylether)-N,N,N’,N’-tetraacetic (EGTA), was used to disrupt the E-cadherin interactions between adjacent MDCK cells as previously (28). Cells were either maintained in complete media or treated with 0.3 M EGTA in DMEM for 1 hour at 37°C and 10% CO_2_. Cell-cell junctional interactions were recovered by replacing the EGTA-containing media with complete media and incubating cells for 1 hour at 37°C and 10% CO_2_. Analysis of pressure, actin – E-cadherin colocalization, and motility was performed as described above.

### Western Blotting

After treatment, MDCKs were lysed using RIPA buffer (1% Triton X-100, 0.5% sodium deoxycholate, 0.1% SDS, 150 mM NaCl, and 50mM Tris, pH8) supplemented with 1X protease inhibitors without EDTA (Roche). Cleared lysates collected were combined with an equal volume of 2X reducing sample buffer, heated to 95°C for 5 min, resolved on 4-12% Tris-glycine polyacrylamide gel (Thermo Fisher Scientific), and then transferred to nitrocellulose membranes containing 0.45 μm pores (Thomas Scientific). Secondary antibodies: goat anti-rabbit IgG Alexa 680 and goat anti-mouse IgG Alexa 680 (Thermo Fisher Scientific). Blots were scanned using Odyssey Scanner (LI-COR Biosciences). Western blots were quantified by normalizing the signal intensity of the E-cadherin or vimentin bands to the corresponding GAPDH signal following background subtraction.

### Statistical methods

Results are presented as the mean ± SEM. One-way analysis of variance with Tukey post-hoc tests were used to compare three or more variables. All comparisons were performed with Prism 7 (GraphPad Software, Inc.). Differences were considered statistically significant at P < 0.05.

## Acknowledgements

We thank Dr. Pavan Vedula and Dr. Tali Gidalevitz for their suggestions on the manuscript. Research reported in this publication was supported by the National Institute of General Medical Sciences of the National Institutes of Health under Award Numbers R01GM126054. There authors declare no competing financial interests.

## Supplemental Material

**Figure S1.**
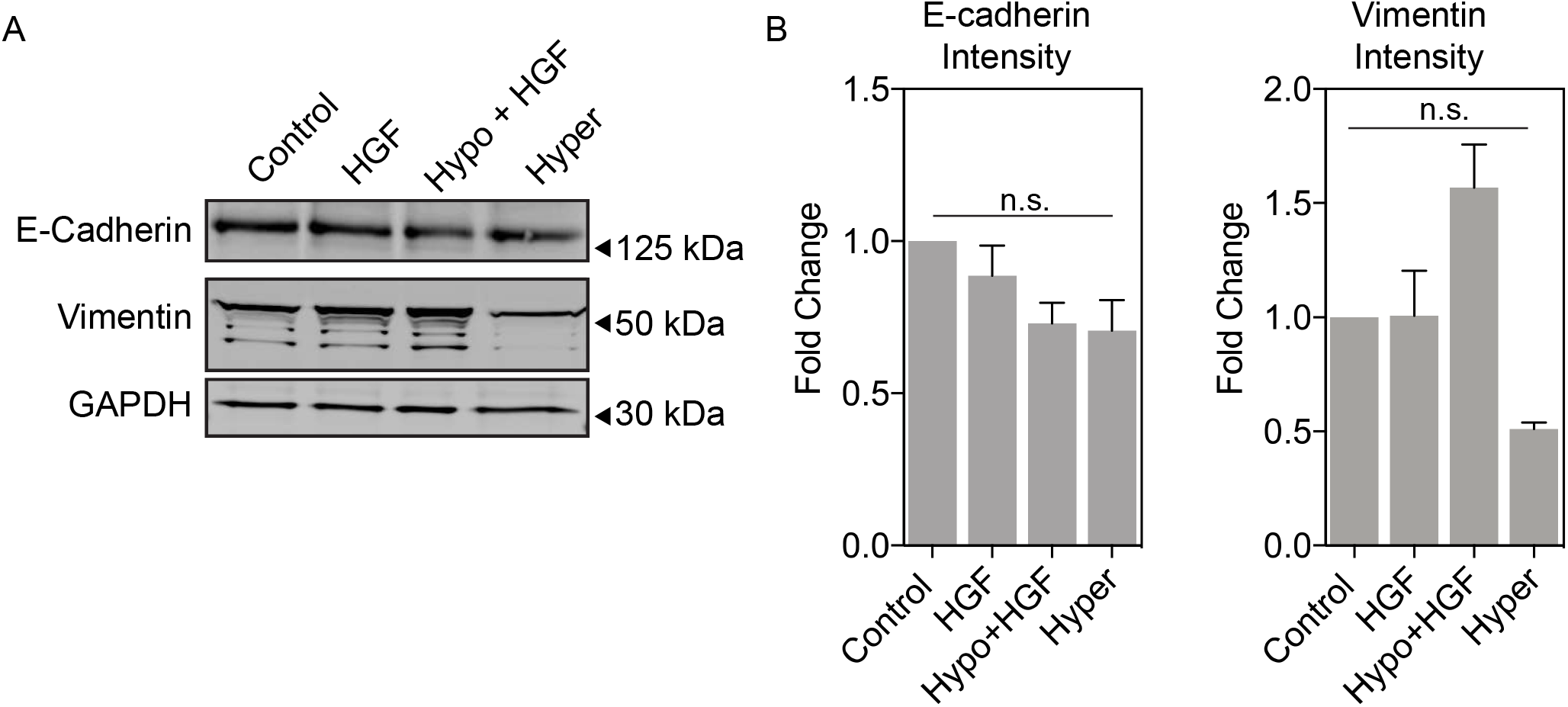
The reduction of cytoplasmic pressure in response to HGF treatment precedes changes in E-cadherin and vimentin expression. (A) A representative western blot of parental MDCK cells after 1 hour of HGF treatment. There was no significant change in E-cadherin or vimentin expression compared to control cells. Vimentin expression did show greater variability in cells treated with a combination of HGF and hypo-osmotic media or hyperosmotic media alone, but the differences were not statistically significant compared to control cells. Quantified in (B) (N = 3). E-cadherin, ns, P > 0.1 versus control. Vimentin, ns, P > 0.07 versus control.

Movie S1. **HGF treatment triggers epithelial cell scattering.** Image sequence of untreated control (left panel) and HGF treated parental MDCK cells (right panel). The increase in movement and the dissociation of cells from each other following exposure to HGF corresponds with the significant drop in pressure in Fig. 2 D.

Movie S2. **Inhibition of NMII is sufficient to trigger epithelial cell scattering.** Image sequence of untreated control (left panel) and blebbistatin treated parental MDCK cells (right panel). Similar to HGF treatment, inhibition of NMII activity triggers loss of cell-cell contacts and increases cell motility, as reported previously (10). This increase in cell migration following NMII inhibition corresponds with the significant drop in cytoplasmic pressure in Fig. 2 E.

Movie S3. **Arp2/3 activity is required for the increase in epithelial cell motility following HGF treatment.** Image sequence of untreated control cells (upper left panel), HGF treated cells (upper right panel), CK666 and CK869 treated cells (lower left panel), and cells treated with a combination of HGF and the Arp2/3 inhibitors (lower right panel). Inhibiting Arp2/3 when treating with HGF significantly reduces cell velocity compared to cells treated with HGF alone. This reduced velocity corresponds with elevated intracellular pressure (Fig. 5 C).

